# Placental immune factors change during the first half of healthy pregnancy

**DOI:** 10.1101/2025.10.16.682794

**Authors:** Maria Obolenska, Olexandr Lykhenko, Yevhenii Kukuruza, Volodymyr Martynenko, Yehor Poliakov, Olga Martsenyuk, Volodymyr Bezguba, Berthold Huppertz

## Abstract

During gestation, the human placenta develops as a fetal organ in direct contact with maternal cells and tissues. Most pregnancy complications, such as preeclampsia and fetal growth restriction, have their roots early in pregnancy, most probably in dysregulation of placental development and/or disturbances in the interaction with maternal (immune) cells. Here, we applied an integrative analysis of open-access gene expression data on the human placenta, comparing expression levels between the first and second trimesters of healthy pregnancy, with a focus on differentially expressed genes related to immune processes. The holistic approach of our study revealed differentially expressed genes involved in the recognition and elimination of potential PAMPs and DAMPs, transendothelial cell migration, cytokine production, transplacental IgG transport, and maintenance of immune tolerance during the transition from the first to the second trimester of placental development. Most DEGs show an increased expression. Only a few genes, ***DEFB1***, ***SLPI***, and ***LCN2,*** which encode antimicrobial proteins, homeostatic chemokines, and a pleiotropic ***PAEP*** that maintains tolerance, display maximal expression in the first trimester, followed by further down-regulation. The cell types deconvolutions revealed the typical representation of placental cells.

## Introduction

The human placenta is a temporary fetal organ that develops into a multifunctional morphological structure, maintaining a close connection between mother and fetus throughout pregnancy and ensuring its successful completion. Disturbances and alterations of placental development and functioning are the cause of pregnancy complications, including fetal growth restriction (FGR) (3-7% of all pregnancies) and preeclampsia (5-10% of all pregnancies^1,2,3^), which can have long-term effects on both the mother and the fetus^4^.

The early-onset subgroup of preeclampsia (high blood pressure or/and proteinuria observed starting around the 20^th^ week of gestation) accounts for about 15-20% of all preeclampsia cases. Due to the lack of a reliable, cost-effective early screening test, well-established primary prevention measures or specific treatment options are a pressing issue^5^.

The etiologies of the early-onset subgroups of FGR and preeclampsia are considered related to adverse placentation and trophoblast invasion, where immune processes and cellular interactions play pivotal roles^1,6^. The maternal organism has to tolerate the persistence of paternal alloantigens without affecting the anti-infectious response against potential pathogens or damaged self-material^7,8^. So far, case-control studies on the human placenta have investigated the expression of single immune-related genes. Other studies have characterized biological processes as part of transcriptome profiling and have identified a number of genes participating in these processes^8,9^.

This study aims to analyze the changes in large-scale gene expression related to immune processes in the human placenta during the critical first half of a healthy pregnancy. To address this challenge, we used gene expression data on the human placenta in health and disease from open-access databases. This data is available for secondary analysis.

We identified 250 differentially expressed genes (DEGs) between the first and second trimesters and 54 DEGs related to the immune processes. We grouped the genes of the immune cluster according to their predominant functions and identified changes in gene expression in the complement system, receptors responsible for recognition and further elimination of pathogen- and damage-associated patterns, activated diapedesis and proinflammatory chemokine and cytokine production, and maintenance of immune tolerance pointing to an activated proinflammatory response. The described changes in gene expression in the human placenta during the transition period from the first to the second trimester may serve as a reference point for investigating the processes in the pathology-affected placenta.

## 2. Methods (see Document S1. Table S1)

Data for the analysis was obtained from the Gene Expression Omnibus (GEO) and BioStudies (ArrayExpress) databases. As a first step, we developed the specialized IGEA (integrative gene expression analysis) database [https://molecularbiology.ink/igea/] containing the metadata for the genome-wide transcriptome profiling of more than 1000 samples of normal and pathology-affected human placentas. The metadata include information on the biological specimen, diagnosis, gestational age, mode of delivery, and other characteristics, which we previously standardized^10,11^.

Methods used in this study are described in Lykhenko et al., 2021a,b^12,13^. In short, raw microarray data was extracted from GEO and BioStudies (ArrayExpress) databases, complemented with standardized metadata from the IGEA database, and integrated into one dataset using the empirical Bayes method for cross-normalization and batch effect removal. The method of integrative analysis overcomes the problem of small sample sizes in experiments, reduces the bias of various processing strategies, and essentially, presents the transcript levels in quantitative relative units^13,14^. To assess the quality of batch effect removal, we employed principal component analysis (PCA) to project the original multi-dimensional data into a two-dimensional space (Fig. 1). Figure 1A depicts the data distribution before batch effect removal. It is apparent that, in terms of gene expression, samples from datasets GSE122214 and GSE9984 differ most both from each other and from the samples of the other two datasets (principal components dim1 and dim2), even though all expression data belong to the first and second trimesters. These datasets introduce the greatest dispersion (dim1 and dim2). The technical variability disappears after batch effect removal (Fig. 1B). By contrast, the difference between first- and second-trimester samples obscured by batch effect (Fig. 1C) became the primary source of variability after its removal (Fig. 1D). Moreover, the gene expression data can also be grouped by fetal sex, as shown along components dim3 and dim4 (Fig. 1E).

**Figure 1:**
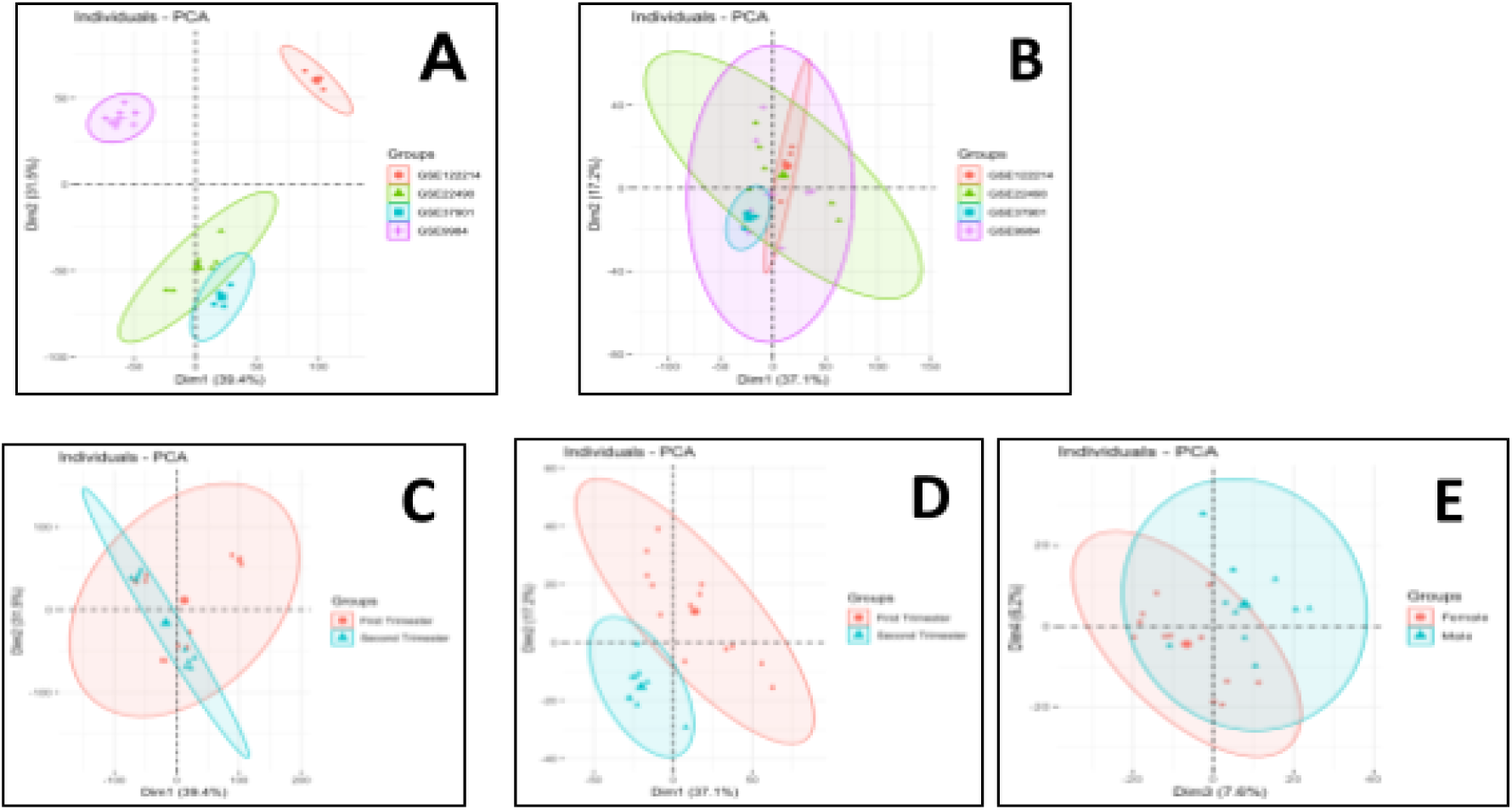
The verification of the batch effect removal during the integration of gene expression data. The PCA plots depict the data distribution before (A and D) and after (B, D and E) batch effect removal.

We used generalized linear models implemented in the R package limma to identify DEGs and the Benjamini**–**Hochberg method to determine the corrected p-value (false discovery rate, FDR) of the difference in gene expression between two time points^15,16^.. For this study, we focused on differences between trimesters. Therefore, we included pregnancy trimester and fetal sex as covariates at the batch effect removal stage to preserve as much biological variation as possible and prevent sex differences overcorrection as batch effects. However, when determining differentially expressed genes in this article, we grouped our expression data by trimester. In our earlier work, we identified moderate autosomal differences *(PUDP, CACFD1, NCAM2, YIPF6, PLXDC1, BTBD1, and GRINA)* in placentas during the first half of pregnancy that correlate with fetus sex^17,18^. We constructed a graph of association and interaction between proteins encoded by DEGs with data from the STRINGdb^19^ using the R STRINGdb package. Then, we clustered and visualized the graph using the fast greedy algorithm from the R igraph package^20^. In the resulting clusters, we identified functionally enriched groups of genes based on the Gene Ontology database using a hypergeometric test, as in Franceschini et al., 2013^21^. The R packages used for the pipeline were obtained from the Bioconductor website bioconductor.org.

Cell type proportions in placental samples were estimated using the nonnegative least squares method of cell-type deconvolution, as implemented in the R MuSiC package^116^, and the placental deconvolution reference developed by Campbell et al. (Seurat object in GSE182381)^117^. Gene Set Enrichment Analysis (GSEA) was performed using the gseGO function from the clusterProfiler R package, focusing on Gene Ontology Biological Process (BP) terms^118^. Genes for GSEA analysis were ranked by π value, a parameter calculated by multiplying logFC and adjusted p-value.

## 3. Results and Discussion

To analyze changes in gene expression levels between the first and second trimesters in healthy placentas, we used samples of villous tissues from 6.4 to 9 weeks of gestation and 13 to 19 weeks. The samples were obtained after uncomplicated elective termination. The authors of the original papers stated that the electively terminated first-trimester pregnancies had no maternal or fetal clinical complications. The mothers were not exposed to drug or steroid hormones and had a record of regular menstruation. They had no clinically confirmed miscarriages in their medical reproductive history. No fetal anomalies were detected for the second-trimester placentas [see Document S1, Table S1, and references therein].

Analysis of gene expression data on the human placenta from the first and second trimesters revealed 250 differentially expressed genes (DEGs). Here, we focused on genes related to immune processes due to the unique immunological complexity of the maternal-fetal interaction during the transition between the first and second trimester of healthy pregnancies [see Document S1, Table S2]. Since the complex features of biological processes are often caused by the combined effects of quantitatively small but numerous changes in gene expression and more substantial but less numerous ones^22^, we selected the DEGs with a fold change of logarithm to base 2 │LogFC│>1.0 and set the threshold for the applied adjusted p-value at <0.05. Focusing on the immune processes, we arranged the genes into functional groups according to their primary previously annotated functions. We decided to analyze only genes that have been proven to play a role in the immune response. However, it is quite probable that other multifunctional proteins indirectly contribute to the reaction to pregnancy as well, They will not be covered in this study, as even the well-established immune pathways are complex enough to warrant further investigation. In the text below, the DEGs, their corresponding proteins, and LogFC values are highlighted in parentheses and bold font format.

### 3.1. Recognition and elimination of potential pathogen-associated and damage-associated molecular patterns

The genes assigned to this group recognize and eliminate potential pathogen-associated and damage-associated molecular patterns, PAMPs and DAMPs, by different mechanisms through host proteins specifically bound to the antigens (opsonin receptors) or by direct binding (non-opsonin receptors).

#### The involvement of the complement system in placental immune response

Our data revealed the involvement of the complement system’s classical, lectin, and alternative pathways in the placental immune response. The ***C1QA*** (**1.37**), ***C1QB*** (**1.33**), and ***C1QC*** (**1.12)** genes encode opsonin receptors that initiate the *classical pathway* (Fig. 2). The respective proteins combine into the C1q complex that is instrumental in recognizing the antigen-antibody immune complexes and initiates autocatalytic activation of the classic complement cascade^23,24^. C1q, as a primary link between innate and acquired immunity^25^, is predominantly detected in the maternal part of the placenta between endovascular trophoblasts invading maternal vessels and decidual endothelial cells^26,27^.

**Figure 2:**
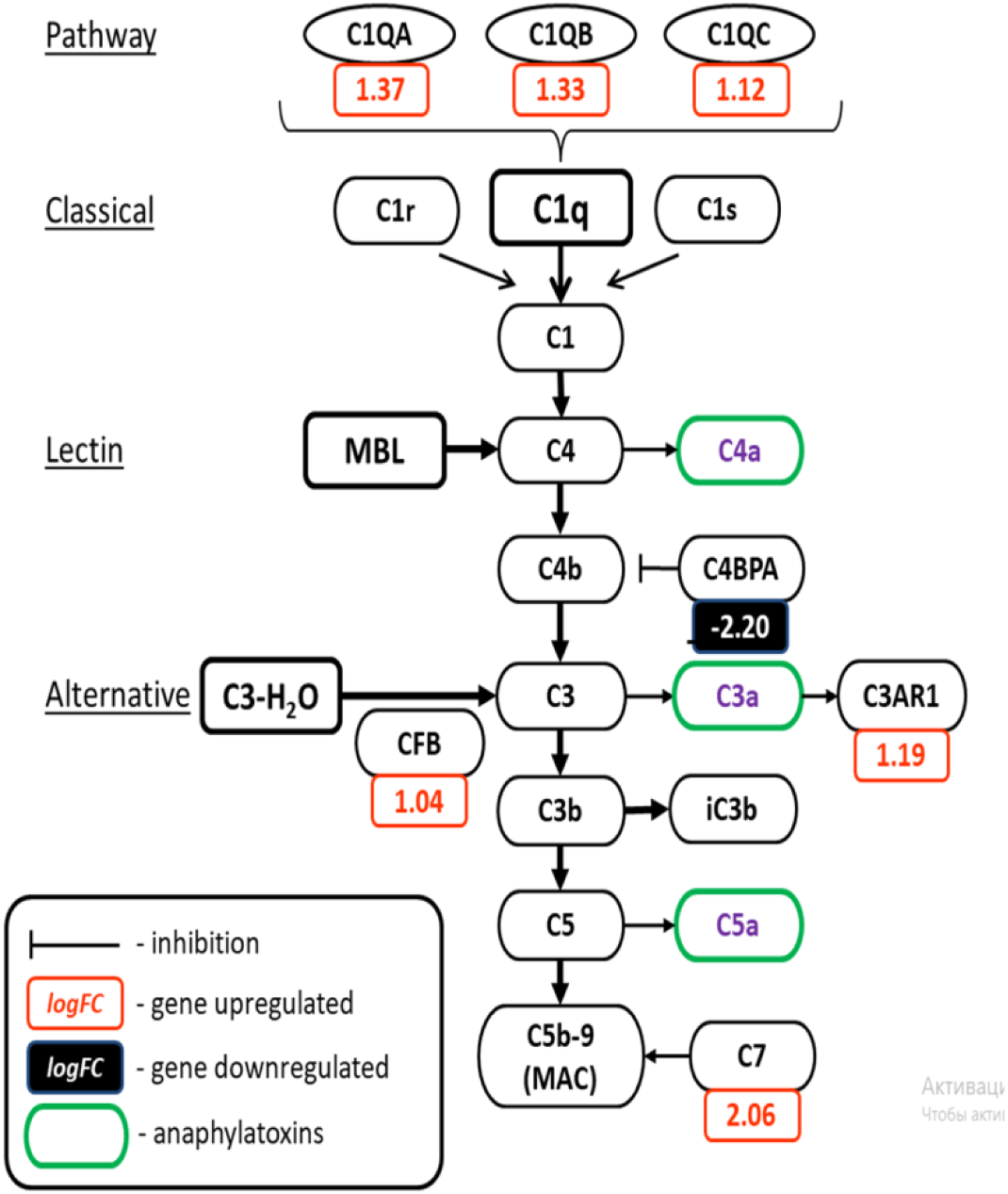
The participation of differentially expressed genes (DEGs) in the complement system. The schematic drawing displays the complement system with its three pathways, the classical pathway via C1, the lectin pathway via MBL, and the alternative pathway via C3-H_2_O. DEGs are shown with the fold change in red (increased expression in the second trimester) or black (decreased expression in the second trimester compared to the first trimester) under the gene name. Anaphylatoxins are marked with a green frame. Schematic drawing adapted from Yednock et al., 2022^32^.

Unlike these proteins, **C4b-binding protein alpha** (***C4BPA,* -2.20**) inhibits the major fragment C4b of the central protein C4 in the *classical* and *lectin pathways*^23^. A decrease in ***C4BPA*** gene expression might reduce (or eliminate) the inhibitory activity of its protein and thus support the further reactions of the complement cascade with the active C4b fragment^28^.

**Complement factor B** (***CFB***, **1.04**) facilitates the catalytic activity of C3 convertase in the *alternative pathway*^29^. C3 convertase cleaves C3 into its active components, C3a and C3b. C3a elicits activity via the **C3A receptor 1 (*C3AR1,* 1.19}** and C3A/**C3AR1** may activate T cells and macrophages and stimulate angiogenesis, chemotaxis, cytokine release, and mast cell degranulation^30,31^.

The **C7 protein** (***C7***, **2.06**) is a component of the MAC complex, which destroys the pathogen’s membrane and the pathogen itself. The elevated expression of ***C7*** aligns with the results of Bulla et al. 2012^27,32^.

#### The recognition receptors during the first to second trimester transition

The scavenger receptors are non-opsonin receptors and directly bind to PAMPs and DAMPs^33^. **Platelet glycoprotein 4 (*CD36,* 1.72)**, for example may recognize various bacteria, viruses, and numerous endogenous ligands such as lipoproteins and HDL (high-density lipoprotein). **CD36** may interact with transmembrane proteins like TLR2, TLR4, and TLR6 in joint pathogen recognition and signal transduction^34,35^. The **CD163** receptor ***(CD163***, **1.29**) is a gram-positive and gram-negative bacteria sensor^36^ and inducer of local inflammation by producing cytokines, e.g., IL-6 and IL-10. In the human placenta, **CD163** was detected in Hofbauer cells. A cleaved soluble form of the **CD163 receptor**, sCD163, was identified in maternal blood as a risk-predictor for developing gestational diabetes mellitus^37^.

The soluble **CD93** receptor (***CD93,* 1.02**) recognizes bacterial DNA rich in unmethylated CpG dinucleotides and delivers it to the endosome. **CD93** is detected in the cytotrophoblast and placental vessels^38^.

The **mannose receptor C-type 1** (***MRC1, 1.17*)** binds mannose-rich structures on the surface of pathogenic viruses, bacteria, and fungi (e.g., Candida albicans) and interacts with glycoproteins released into the circulation during the inflammatory response.

The **FPR1** receptor (***FPR1,* 1.43**) interacts with N-formyl-methionine-peptides, by which degraded bacteria and host cells’ damaged mitochondria signal to immune cells: “Find me, Catch me, and Eat me.” The timely reaction to these signals transmitted via **FPR1** promotes phagocytosis preventing the destruction of the damaged cell’s plasma membrane, hydrolytic enzyme leakage, and the development of autoimmunity^39^.

Two genes encode for **G-protein coupled receptors**. **GPR34** (**1.55**) is located on the X-chromosome and encodes the orphan **receptor.** The gene ***GPR183*** (**1.54**) is tissue-specifically highly expressed by immune cells in the human placenta and brain. The immune cells are driven by oxysterol gradient to locations of high ligand concentration^40^.

All structures bound to recognition receptors mentioned above are eliminated by phagocytosis. The toll-like receptor 7 (***TLR7***, 1.4) is located in the endosome; it recognizes ssRNA (viral) RNA and transmits a signal to the nucleus via transcription factor IRF (interferon regulatory factor). IRF, in turn, induces transcription of the IFNα/β and type III IFN genes, and an antiviral response^41,42^. **TLR7** is detected in first-trimester cytotrophoblast and second-trimester Hofbauer cells^43^.

#### Elimination of potential pathogens and damaged cells’ material

Most recognition receptors trigger the internalization of particles to their destruction in the phagolysosome where NADPH oxidase (NOX)-derived reactive oxygen species (ROS) provide antimicrobial activity^44,45,46^. The formation of NOX is initiated by phagocytosis via the assembly of six subunits, among which are **cytochrome b-245 heavy chain (*CYBB,* 1.47), neutrophil cytosol factor 2 (*NCF2,* 1.10**), and **neutrophil cytosol factor 4 (*NCF4,* 1.03**). NOX shuttles electrons from cytoplasmic NADPH to molecular oxygen in phagosomes and the extracellular space to produce oxidants (O_2_−, HO•, and H_2_O_2_) with bactericidal activity. The ROS might also be implicated in signal transduction and regulation of proliferation, differentiation, migration, and cell death^44,47^. NOX as the primary source of superoxide in first trimester chorionic villi may be required for redox adaptation to increased oxygen pressure after the 12^th^ week of gestation when hemotrophic nutrition is established^48,49^.

Upregulated ***RNase 6*** (**1.13**) encodes an enzyme that acts against gram-positive and gram-negative bacteria^50,51^. In humans, ***RNase 6*** expression is mainly localized in the placenta and the urinary tract during urinary tract infections^52,53^. **RNase 6** acts as a chemoattractant, damage-associated molecule or alarmin, immune cell activator, and opsonin^50,51^.

The ***BST2*** gene (**1.01**) encodes **tetherin**, which retains virions in infected cells near the cell membrane to prevent their spread inside and outside the cell. The **BST2 protein** can activate interferon-stimulated genes in response to infection by enveloping viruses^54^.

However, not all “protective” bactericidal genes are upregulated during this period. The level of beta-defensin 1 (*DEFB1,* -3.23), secretory leukocyte protease inhibitor (*SLPI,*-2.18), and **neutrophil gelatinase-associated lipocalin**, NGAL (*LCN2,* -1.98) might be highest in the first trimester and decline later. **DEFB 1** is mainly expressed in epithelial and immune cells of the reproductive tract^55,56^. **DEFB1** may directly inactivate and inhibit the replication of various viruses, kill bacteria, or inhibit their growth^57^. *DEFB1* expression has been detected in the chorion, decidua, and amnion^58^. Peak endometrial mRNA expression of the *DEFB1* gene coincides with implantation and menstruation, suggesting its protective function during these critical events^58^.

Like DEFB1, **SLPI** is a natural antimicrobial of the innate immune response. **SLPI** guards the host from infection and maintains homeostasis^59,60,61^ while excessive **SLPI** expression increases epithelial tumors’ metastatic potential^61^. In the human placenta, **SLPI** has been detected in the decidua and amnion^58^, where it may limit proteinase activities during the implantation period and regulate the extent of trophoblast invasion^62^.

**NGAL** is an active bacteriostatic that deprives prokaryotic cells of iron, thus delaying their growth and reproduction. **NGAL** is a neutrophil attractant, involved in apoptosis, innate immunity, and kidney development, and found in neutrophil and macrophage granules^63^.

The high starting expression levels of these three genes might be the result of the protective innate immune response to the destruction of the endometrium during implantation. In contrast, the subsequent reduced expression concords with the increasing expression of pro-inflammatory factors.

### 3.2 Immune cell migration, cytokines, and immune tolerance (see Document S1, Table S3)

#### Transendothelial immune cell migration (*Fig. 3*)

Several DEGs are directly or indirectly involved in diapedesis, cell transmigration from the bloodstream into the surrounding tissue^64,65^. Histamine is a potent vasodilator that aids in the diapedesis of leukocytes^66^. **Diamine oxidase (DAO) (*AOC1,* 1.65**) is the only extracellular enzyme capable of inactivating histamine in humans and preventing its accumulation in the placenta, thus decreasing the risk of premature uterine contractions and fetal death^67^.

**Figure 3:**
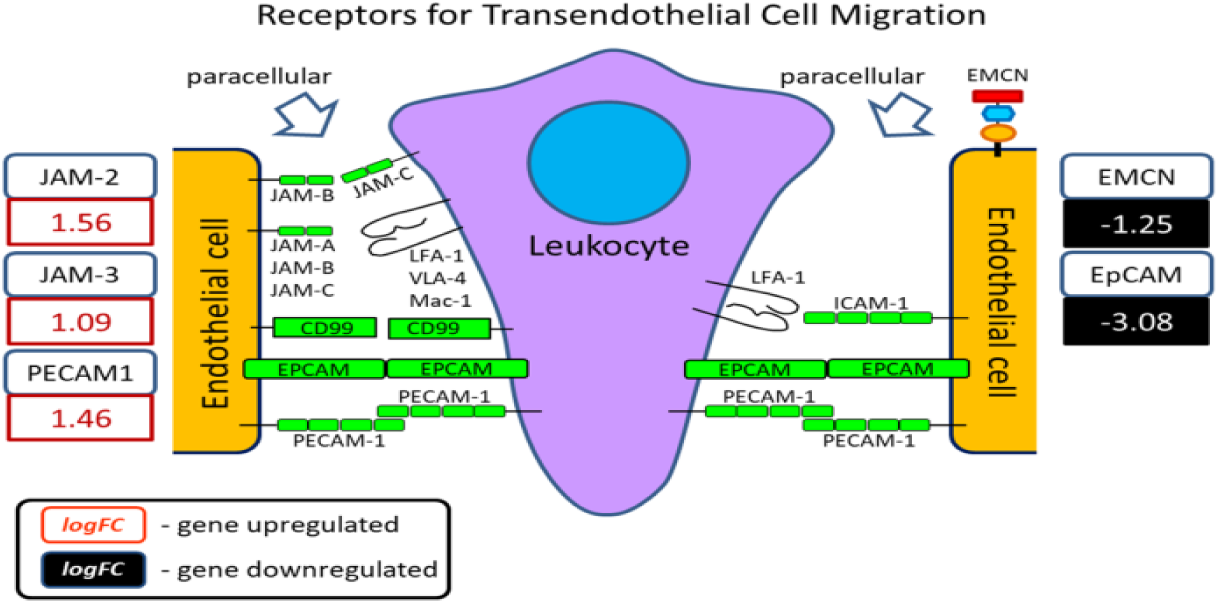
Receptors involved in regulating transendothelial cell migration. The schematic drawing shows the migration of a leukocyte (violet, center) between two endothelial cells (orange, margin). Differentially expressed genes (DEGs) are shown with the fold change in red (increased expression in the second trimester) or black (decreased expression in the second trimester compared to the first trimester) under the gene name. Schematic drawing adapted from Strell & Entschladen, 2008^125^.

The **platelet endothelial cell adhesion molecule 1** (***PECAM1,* 1.46**) and **junctional adhesion molecules B** (***JAM-2,* 1.56**) and **C** (***JAM-3,* 1.09**) belong to the immunoglobulin superfamily and are expressed by leukocytes, platelets, and endothelial cells. Endothelial **PECAM1** interacts with leukocyte **PECAM1** and supports leukocyte diapedesis by dephosphorylation of VE-cadherin^68^. **JAM proteins** are located in tight junctions between endothelial cells. **JAM-B** binds to α4β1 integrin and **JAM-C** to αMβ2 integrin on leukocytes. Endothelial **JAM-B** can connect with leukocyte **JAM-C**^69^. The concentration of these proteins in the human placental villous cytotrophoblast and endothelium is ten times higher than the average concentration in other tissues [Human Protein Atlas, HPA] thus enhancing the diapedesis.

Upregulated ***Nectin-3*** (**1.48**) plays a significant part in facilitating the movement of T-lymphocytes and monocytes across the endothelial barrier^70^.

Two other proteins, **endomucin** (***EMCN,*** -**1.25**) and epithelial cell adhesion molecule (*EpCAM,* -3.08), might decline from the first to the second trimester. Endomucin is mainly synthesized by endothelial cells of postcapillary venules, where it prevents leukocyte adhesion and diapedesis. Reduced expression of the ***EMCN*** gene suggests a potential increase in the number of leukocyte attachments and diapedesis during this period. **EpCAM** on the surface of an epithelial cell binds to **EpCAM** on the surface of a neighboring cell, keeping the cells together. The decline in *EpCAM* expression suggests an increase in the facilitation of diapedesis through the placental barrier. The maximum of **EpCAM** expression level was detected in 4–5 week placentas. After that, it quickly diminished in the late first trimester^71^, as our integrative analysis confirmed.

#### Chemokines in cell migration (*Fig. 4*)

The migration of immune cells from the bloodstream into the surrounding tissue is regulated by chemokines, a large family of small, secreted proteins that transmit signals through cell surface G protein-coupled chemokine receptors. The differentially expressed genes encoding chemokines demonstrate remarkable synchronicity. The genes for homeostatic chemokines, ***CCL21*** (**-1.24**), ***CXCL12*** (**-1.17**), ***CXCL13*** (**-2.18**), and ***CCR7* receptor** (**-1.26**) are downregulated, while the genes for pro-inflammatory chemokines ***CCL2*** *(**1.6*****5**), ***CCL8*** (**1.14**), ***CCL13*** (**1.51**), ***CXCL1*** (**1.53**), ***CXCL2*** (**1.37**), ***CXCL4*** (**1.98**), ***CXCL7*** (**2.0**), and ***CXCL8*** (i.e. ***IL8***) (**1.74**) are upregulated. Their function is not limited by chemotaxis. The homeostatic **CCL21** and **CCR7** interaction promotes epithelial-to-mesenchymal transition (EMT) in a trophoblast-derived cell line^72^. The level of **CXCL12** expression in all trophoblast subtypes is closely related to the invasive depth of trophoblast cells. The placental **CXCL12** may promote Th2 immunity^73,74^. ***СXCL13*** is expressed in decidual stromal cells, fetal membranes, and the umbilical vein, where it may recruit T and B cells and regulate the adaptive immune response. The homeostatic genes ***CXCL12***, ***CXCL13***, ***CCL21,*** and ***CCR7* receptor** contain the hypoxia response element (HSE) in their promoters. They might be the targets of heterodimeric transcription factor HIF-1a–HIF-1b, which is activated by hypoxia typical for the human placenta before the 10^th^/12^th^ week of gestation and the onset of the maternal blood supply^119,120^.

**Figure 4:**
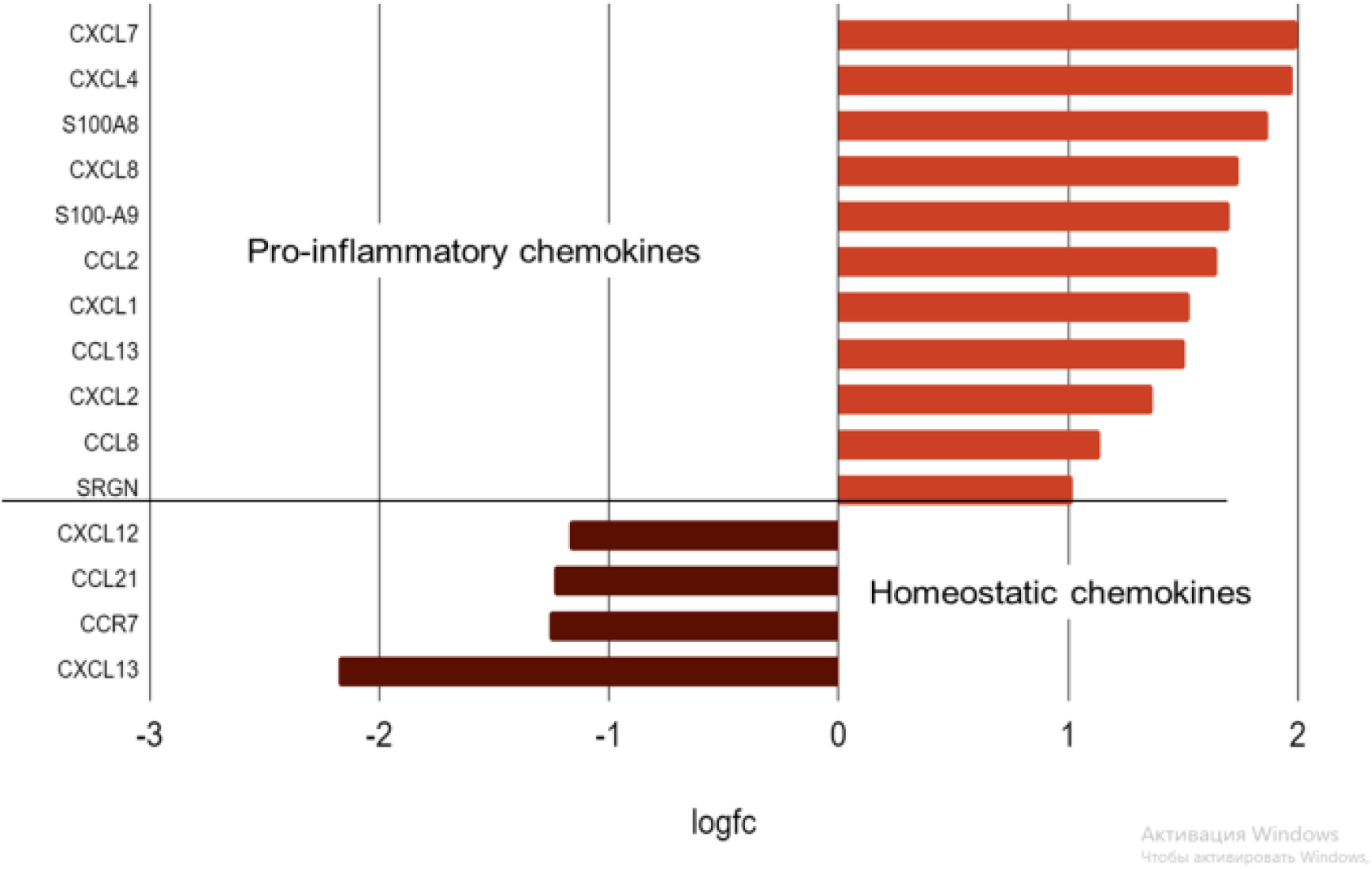
Changes in the expression of genes encoding homeostatic and pro-inflammatory chemokines in the human placenta between the first and second trimesters of healthy pregnancy. During the second trimester, the expression of the pro-inflammatory chemokines increases, while the expression of the homeostatic chemokines is downregulated.

**Figure 5:**
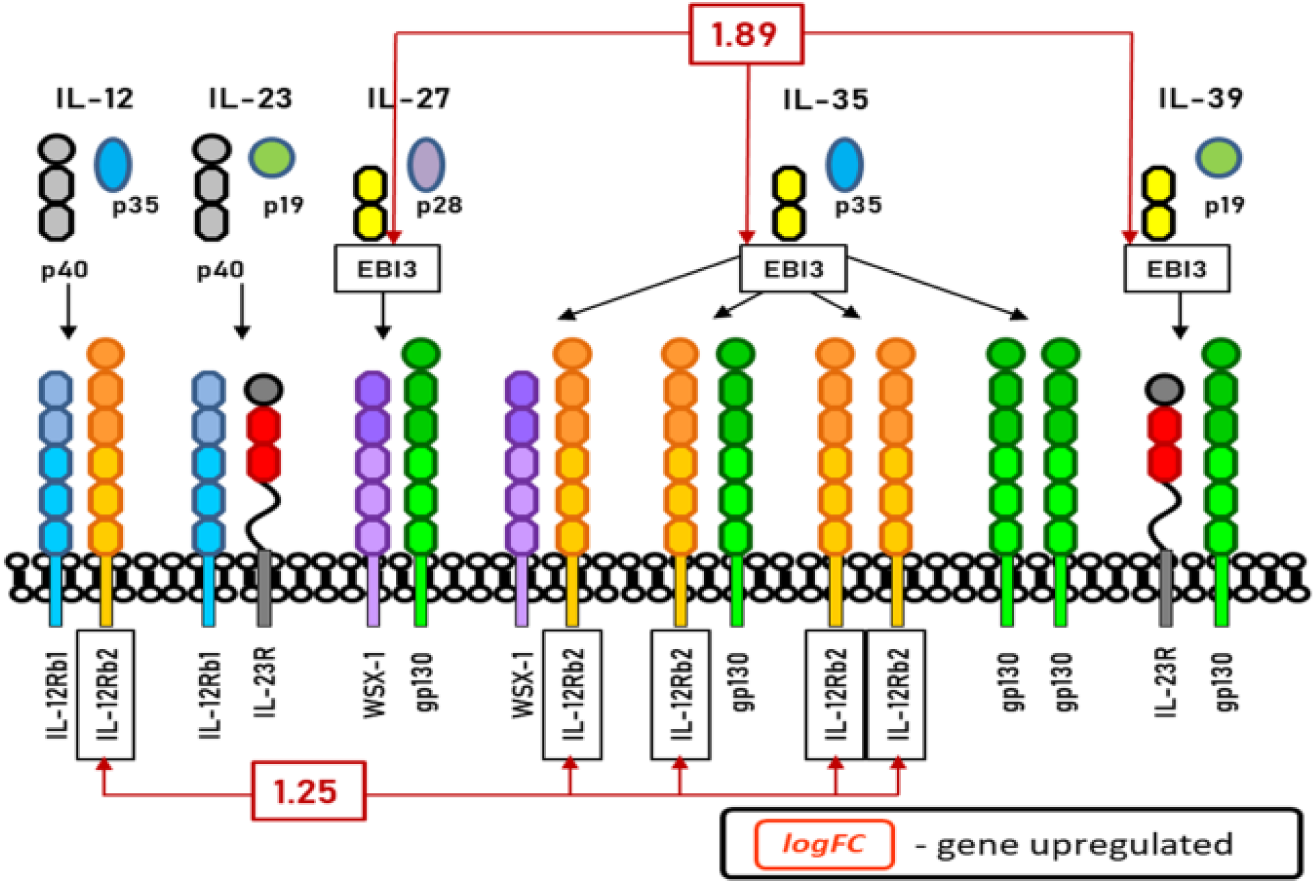
Differentially expressed genes of the IL-6/IL-12 family. The IL6/IL12 cytokine family displays a multitude of different receptor and ligand pairs and interactions. Two differentially expressed genes in combinatorial ligand/receptor complexes of the IL6/IL12 cytokine family were identified. Schematic drawing adapted from Floss et al., 2020.^99^

The gradual decline of homeostatic gene expression appears to give way to inflammatory chemokines usually formed under potential pro-inflammatory stimuli of Il-1, TNF-α, LPS, and viruses^75^. Inflammatory chemokines attract immune cells to inflammatory sites and induce the release of superoxide ions, acting in inflammation, proliferation, metabolism, and angiogenesis^75,76,77^. **Interleukin 8** (chemokine **CXCL8)** is a major pro-inflammatory cytokine. It was localized in villous trophoblast and Hofbauer cells, decidual stromal cells, decidual lymphocytes, and endometrial gland cells^78^. Unlike many other chemokines, **IL-8** has distinct target specificity for neutrophils, with weak effects on other blood cells^79^. It enhances the oxidative metabolism and generation of ROS, possibly leading to oxidative stress^79,80^. Studies on placental villous explants from the first and second trimesters show that **IL-8** secretion increases during gestation, with maximal production in the second trimester and at term^81^. Thus, on ***IL-8*** mRNA level, our data confirms and complements the previously obtained results.

**CCL2**, localized in trophoblast and decidual cells, may attract macrophages, T cells, basophils, mast cells, and NK cells. These cells assemble at the maternal-fetal interface, guaranteeing the immune microenvironment and tolerance^82^.

***CXCL4*** and ***CXCL7*** mRNAs originate from maternal platelets adhering to the surface of the placental syncytiotrophoblast^83^. Their corresponding proteins, **platelet factor 4** (**PF4**) and **pro-platelet basic protein** (**PPBP**), are released upon platelet activation. **PF4** promotes fibrinoid deposition on the syncytiotrophoblast^84,85^, interacts with the upregulated **blood coagulation factor XIIIA** (***F13A1,* 1.29**), and regulates trophoblast invasion (by activating CCR1 and the **macrophage inflammatory protein-1** (***CCL3,* 1.52** ) ^83,86^. The protein **PPBP** has been shown to influence the synthesis of DNA, hyaluronic acid and sulfated glycosaminoglycan, prostaglandin E2 secretion, and impact glycolysis.

The genes ***S100A8*** (**1.87**) and ***S100A9*** (**1.70**) encode proteins constitutively expressed in neutrophils and monocytes. In response to inflammatory stimuli, **S100A8** and **S100A9** form a complex (calprotectin), which interacts with TLR4, RAGE, and CD33 receptors and acts as an alarmin or DAMP. Calprotectin may induce neutrophil degranulation with bactericidal effects, production and release of pro-inflammatory chemokines, cytokines, and ROS, leukocyte recruitment to the inflammatory site, generating a positive, pro-inflammatory feedback loop^87,88^.

The proteoglycan **serglycin (*SRGN,* 1.02**) is considered a chemokine as it interacts with numerous pro-inflammatory mediators and regulates their retention and secretion from secretory granules^89^.

#### The IL-6/IL-12 and the CSF cytokine families with shared ligand/receptor elements

The cytokine families IL-6/IL-12 and CSF play a pivotal role in pregnancy-related processes. The combination of common and specific components in the heteromeric structures of these cytokines and their receptors ensures the variability of cytokine/receptor complexes with distinct biological functions. The IL-6/IL-12 family includes IL-6, IL-12, IL-23, IL-27, IL-35, and IL-39 cytokines. **Interleukin-6** (***IL-6,* 1.69**) is a major driver of physiological and pathological pregnancy-related processes^79^. It is implicated in organ development, acute-phase response, immune response, and metabolic regulation. Elevated levels of **IL-6** point to inflammatory response^79^. In the human placenta, **IL-6**^90^ and IL-6R^91^ were detected in the syncytiotrophoblast and extravillous trophoblast, suggesting trophoblast response for IL-6/IL-6R interaction. **IL-6** production begins as early as late 6^th^ week of gestation, with a subsequent increase until term^92^, which is partially confirmed by our data on the mRNA level increase from the first to the second trimester.

The heterodimeric receptors of *IL-6/IL-12* family cytokines share the common glycoprotein 130 signaling subunit and **interleukin 12 receptor, beta 2 subunit** (***IL-12RВ2,* 1.25**) ^93^. In contrast, IL-27, IL-35, and IL-39 cytokines share the **EBI3** subunit encoded by Epstein-Barr virus-induced gene 3 (***EBI3,* 1.89),** which is expressed by the syncytiotrophoblast and extravillous trophoblast throughout pregnancy^94,,99^. The binding of *IL-6/IL-12* cytokines to their receptors activates JAK-STAT signaling pathways^93,95^.

Similar to the receptors of *IL-6/IL-12* family cytokines, the receptors of colony-stimulating factors share the **β signaling subunit (*CSF2RB,* 1.44**) between GM-CSF, IL-3, and IL-5 receptors^96^. Each cytokine binds to a cytokine-specific α chain of a heterodimeric receptor and recruits the primary signaling **CSF2RB** chain that connects the receptor to downstream signaling^95,96^ (Figure 6). The **CSF2RB** protein was detected in the human placenta, particularly in the cells of primitive hematopoiesis in the early placenta^97,98^.

**Figure 6:**
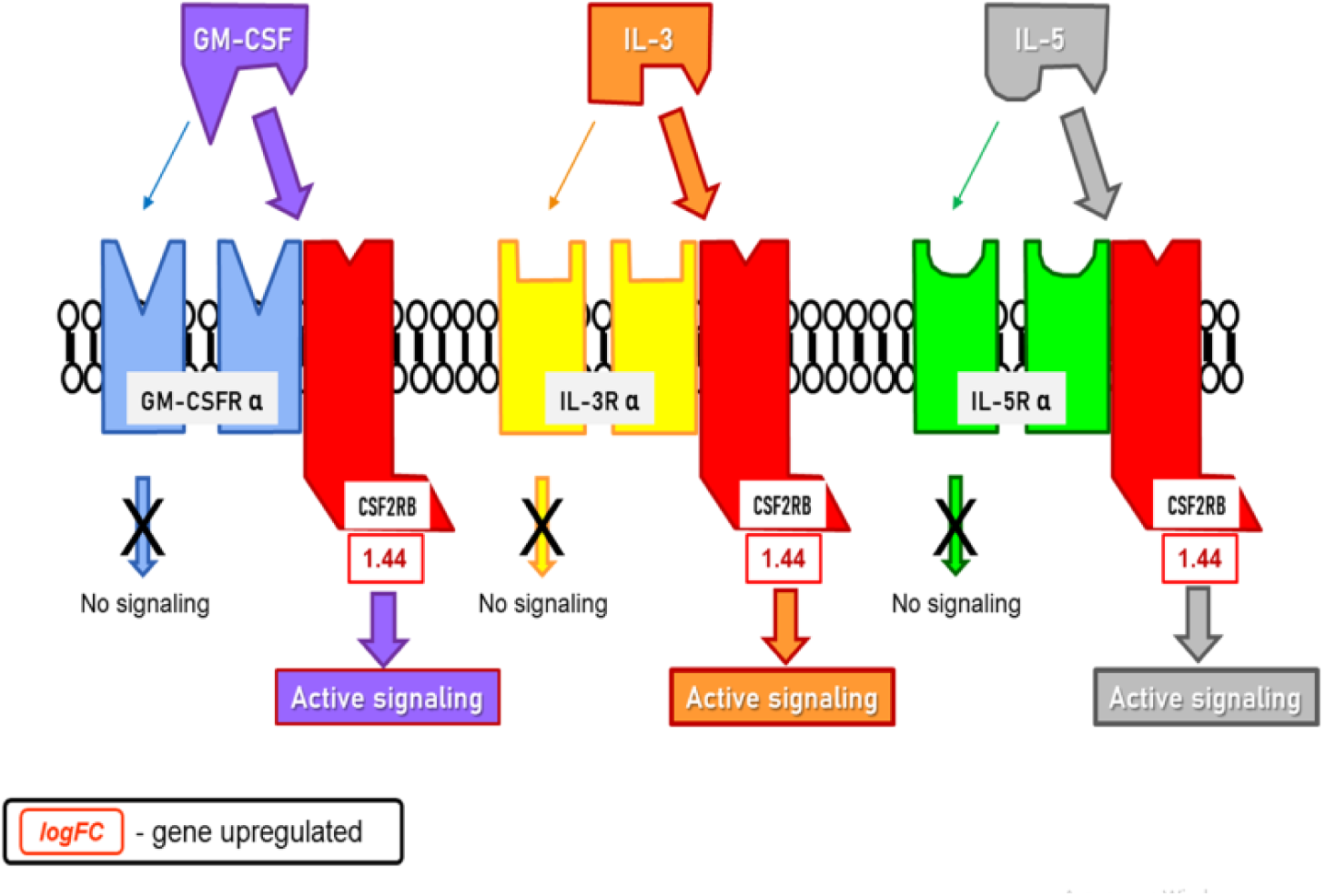
The differentially expressed gene *CSF2RB* in combinatorial ligand/receptor complexes of the CSF cytokine family. The β common ([βc]/CD131) family of cytokines uses different alpha-subunits of receptors with a common beta subunit (CSF2RB). This differentially expressed gene (CSF2RB) facilitates active signaling in combinatorial ligand/receptor complexes of the CSF cytokine family. Schematic drawing adapted from https://www.creative-diagnostics.com/common-beta-chain-receptor-family.htm.

#### Transplacental transport of IgG and immune tolerance

The FCGR2B (***FCGR2B,* 3.0**), a receptor for the Fc fragment of IgGs, plays a crucial role in the transplacental transfer of maternal IgGs to fetal blood. This process, which commences around the 13^th^ week of gestation and peaks at the end of the third trimester, is instrumental in providing the fetus and newborn with passive protection against infectious diseases, especially when their humoral response is still developing^100^.

The **FCGR2B** protein also functions as a negative regulator of immune cell activity, preventing excessive immune reactions and maintaining placental immune tolerance^101^. This activity might be supported by the leukocyte-associated immunoglobulin-like receptor 1 ***(LAIR1,* 1.34)** and leukocyte immunoglobulin like receptor B5 ***(LILRB5, 1.27),*** which maintain placental immune tolerance by a mechanism similar to that of **FCGR2B**^102,103^. **LILRB5** may recognize self-molecules represented by histocompatibility complexes of class I (HLA Class I) and inhibit the various functions of effector immune cells like migration, proliferation, synthesis and secretion of cytokines, and phagocytosis^103^. ***FCGR2B*** is expressed in the endothelium of villous vessels^104^ while ***LAIR1*** — by Hofbauer cells, dendritic cells, macrophages, monocytes, granulocytes, and NK cells [HPA].

The other protein, the **progestagen-associated endometrial protein** or glycodelin (***PAEP,* -4.43**), also plays a substantial role in fetomaternal tolerance, particularly during the earliest gestational period. It regulates blastocyst implantation, trophoblast differentiation, and placental development^105^ The **PAEP**’s isoform glycodelin-A is abundant in the secretory endometrium, decidua, and amniotic fluid^106^. It is secreted from the endometrial glands starting right after the implantation, peaking between 6 and 12 weeks of gestation, and declining after that^106^. Our analysis of the changes in gene expression level is in line with these observations.

### 3.3 Placental cell type deconvolution (Fig. 7, see Document S1, Table S4)

The bulk placental tissue-level gene expression measurements represent a convolution of gene expression signals from individual cells and cell types and depend on their ratios. Although we accompanied most genes’ descriptions with cellular localization information, we were unaware of cell-type proportions. The application of the nonnegative least squares (NNLS) method revealed that fetal fibroblasts constitute 26% to 42% of cells, fetal mesenchymal stem cells — 20% to 29%, and proliferative cytotrophoblasts – invariably 23-25% across placental samples. The lower percentage of cytotrophoblasts (∼15.7%) compared to proliferative ones, and the extremely low amount of syncytiotrophoblasts (1% to 3%) and nucleated red blood cells, align with the results of other studies^121^.

**Figure 7:**
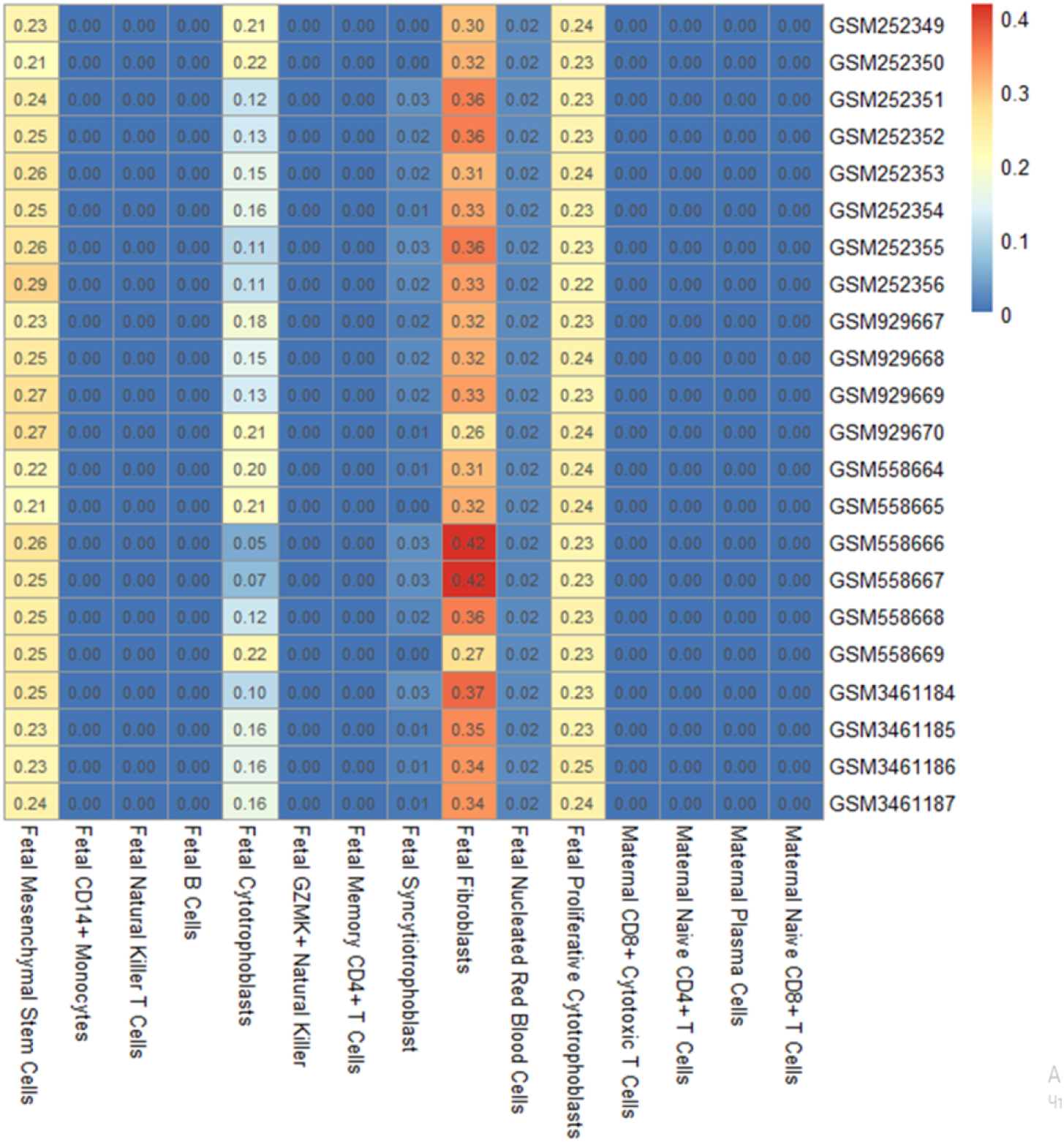
Cell-type proportions as estimated based on nonnegative least squares method of cell-type deconvolution, using the reference dataset developed by Campbell et al^117^. The color scale represents the percentage of each cell type in a sample.

**Figure 8:**
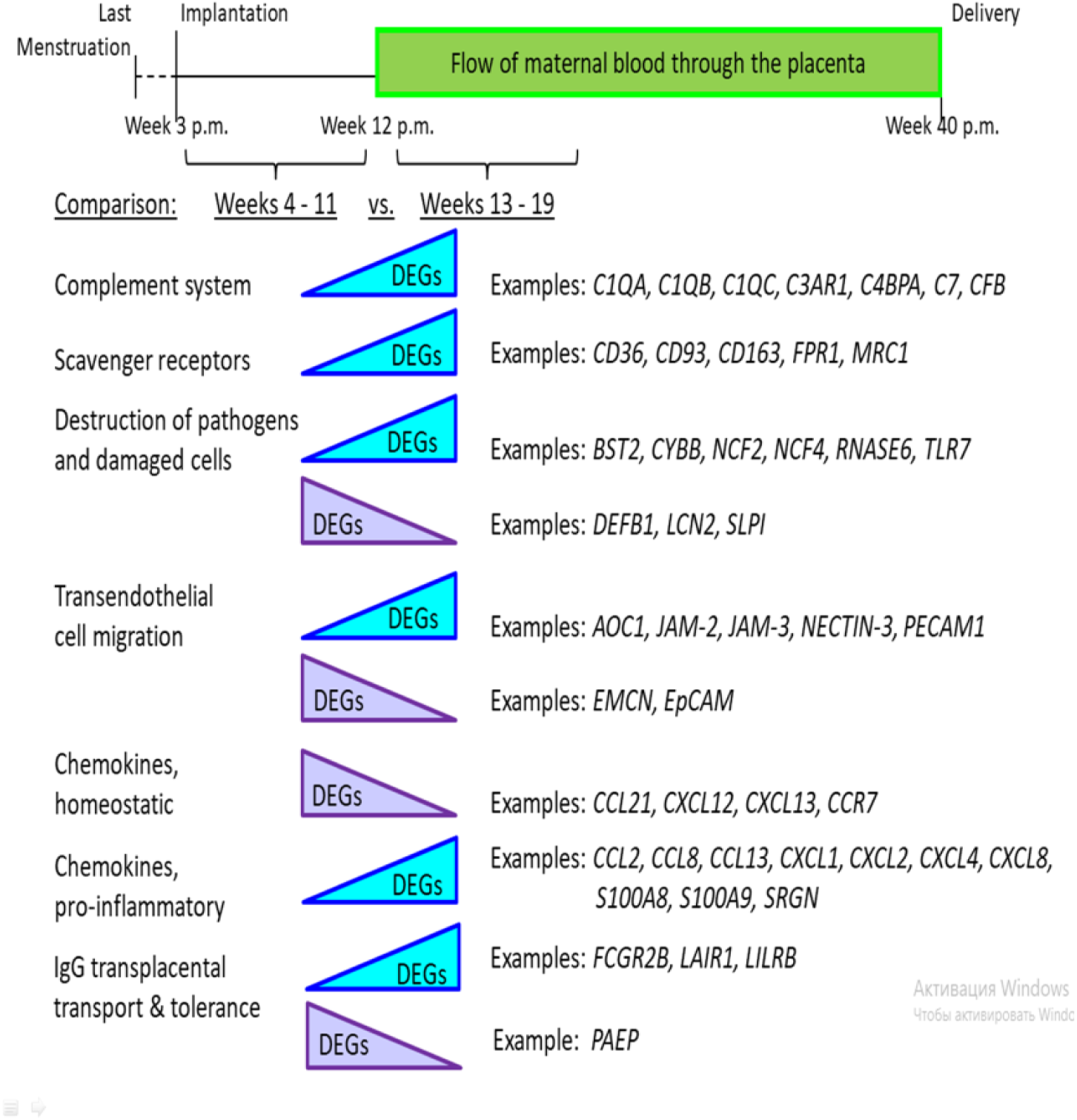
Overview of the changes in immune-related gene expression of the human placenta during the first half of pregnancy. The blue triangles represent an increased and the purple triangles a decreased expression level during the transition from the first to the second trimester of healthy pregnancy. All DEGs listed as examples have been quantified in this study. On top of the schematic drawing, the duration of pregnancy with the last menstrual cycle, implantation, onset of maternal blood flow through the placenta as well as delivery are depicted.

According to Campbell et al.^117^, proliferative cytotrophoblasts are distinguished from other cytotrophoblasts by overexpression of genes related to cytoplasmic translation and mitotic sister chromatin segregation, indicative of their proliferative phenotype. The GSEA analysis applied to differentially expressed genes between the first and second trimesters revealed the highly active cellular proliferation enriched in cell cycle and division processes, chromosome segregation, sister chromatid segregation, nuclear division, and metabolic activity. It is visualized using a treeplot based on semantic similarity (pairwise_termsim) with the enrichplot package (Document S1, Table S4). Thus, it is very likely that the proliferative cytotrophoblasts are responsible for the cellular mitotic activity; the lack of maternal cells in the estimate further suggests that that the analyzed samples are from placenta tissue and not the surrounding maternal tissue.

Summary. The holistic approach of our study revealed differentially expressed genes involved in the recognition and elimination of potential PAMPs and DAMPs, transendothelial cell migration, cytokine production, transplacental IgG transport, and maintenance of immune tolerance during the transition from the first to the second trimester of placental development. Most DEGs show an increased expression. Only a few genes, ***DEFB1***, ***SLPI***, and ***LCN2,*** which encode antimicrobial proteins, homeostatic chemokines, and a pleiotropic ***PAEP*** that maintains tolerance, display maximal expression in the first trimester, followed by further downregulation.

This data aligns with the general concept of immune activation, tolerance maintenance, active proliferation, and morphogenetic processes during early pregnancy^110^. The idea is supported by data from numerous studies, including those that explore transcriptome dynamics using microarray technology and RNA sequencing. For example, comparing our results (four studies) with those of the other three independent studies, which also explored gene expression in placental tissue from the first trimester to midgestation, we observed immune and inflammatory responses, an enriched category of antiviral immunity, extracellular matrix remodeling, responses to cytokines, cell adhesion, and signal transduction activity^111,112,113^. Overall, between 6–10 weeks’ and 11–23 weeks’ gestation, RNAseq profiling of transcriptional dynamics at an individual transcript resolution identified similar changes, together with cell migration, membrane, and cell adhesion processes, cytokine-related signaling,^114,115,122^. Compared to microarray technology, RNA sequencing provides more detailed information about the transcriptome. Along with classical differential gene expression (DEGs), RNA sequencing enables the characterization of transcriptome profiling through differential transcript (isoform) expression (DTE) and differential transcript usage (DTU), which captures changes in transcript proportions^113^. For instance, RNA sequencing has shown that ***CD36* (1.72)** has six isoforms, with only one of them being responsible for the upregulation during early gestation^113^. Precise knowledge of DTU is helpful for a better understanding of molecular processes. Though RNA sequencing has a wider dynamic range, the expression of *the **PAEP*** gene decreases equally between the first and second trimesters, with logFC values of -4.43 and **-4.33,** respectively^114^, indicating the relevance of both methods in detecting principal parameters.

Significant downregulation of ***DEFB1***, ***SLPI***, ***LCN2,*** homeostatic chemokines, and ***PAEP*** is preceded by their elevated expression during the earliest physiological state of low oxygen in placental development. The genes for homeostatic cytokines and ***DEFB1*** contain a response element HRE in their promoters, and they are more likely to be regulated by the HIF1α-HIF1b transcription factor, which is active during this period^123^. *LCN2* is capable of orchestrating the HIF1α pathway^124^. The allocation of a separate stage during the first trimester is consonant with a demarcation of three distinct temporal phases in early placenta, detected by RNA sequencing: early gestation (6-10 weeks), a transition phase (11-13 weeks) and early second trimester to mid gestation (14-23 weeks), with chracterisitc RNA prorfiles for each phase^114^. The maternal blood flow to the placenta between the 10th and 12th weeks triggers a significant change in gene expression, as observed in our studies and others, manifesting as a pronounced proinflammatory and tolerogenic response to initial trophoblast invasion and ongoing uterine remodeling, as well as the recognition of fetal antigens by maternal immune cells. As demonstrated in our study, the activated expression of alarmins or DAMPs, including **S100A8, S100A9**, and **RNASE6,** may play a crucial role in initiating and maintaining a proinflammatory response. The data on gene expression were accompanied by localization of their mRNA and corresponding proteins in placental cells when possible. The described changes in immune-related gene expression in the human placenta during the transition period from the first to the second trimester provide only an indication of potential protein levels and function due to differential translation and posttranslational modifications. Nonetheless, changes in their gene expression are essential constituents within major physiological transformations in the human placenta.

The study’s strengths. This study focuses on a critical period of pregnancy during which many pathologies are believed to originate. Placental gene expression during the transition from the first to the second trimester remains understudied compared to the term placenta, primarily because biological samples, particularly those from the second trimester, are challenging to obtain. Addressing this topic is still relevant. The inquiry into the microarray data in open-access sources enabled us to summarize and interpret the valuable accumulated data. Focusing on the immune processes, we analyzed the differential expression of genes within previously annotated functional groups that had been poorly reported in detail in other studies. We allocated a distinct temporal phase in early pregnancy characterized by a low oxygen state, with corresponding immune–related genes predominantly regulated by HIF1α. We highlighted the special molecular interactions within functional groups, enabling these groups to achieve their optimal operating effectiveness. In the complement system, the activation of effector gene expression occurs alongside the downregulation of the inhibitor gene, thus promoting the efficiency of the complement system. An impressive example of synchronicity and harmonization reveals the expression of chemokine genes. The homeotic chemokines regulated by HIF1α are synchronously downregulated, leading to the following synchronous upregulation of proinflammatory chemokines. The regulatory mechanisms of coexpression of chemokine genes, similar to those regulating other immune-related genes and the cell set in this study, pose a further challenge.

The study’s limitations. As NGS and its various applications, such as single-cell technology and spatial transcriptomics, become increasingly popular, it appears that microarrays are already becoming obsolete. RNA sequencing provides comprehensive coverage of all RNA sequences present in a sample, offering a wider dynamic range and heightened sensitivity that make it the preferred choice for identifying novel transcripts and unraveling complex gene expression patterns. Unlike RNA sequencing, microarray technology detects predefined sequences present on the platform and utilizes hybridization as its primary method for detecting RNA abundance. The data from RNA sequencing is presented in absolute values, while that of microarray is in relative values. Compared with RNA-Seq, the RNA microarray is simpler in both experimental and data analysis processes. The choice of method depends on the goal of the biological research and the financial and instrumental capabilities. Here, we had no choice as we wanted to summarize the valuable data obtained by microarray technology and identify the *most pronounced* functional changes spanning the periods between the first and second trimesters, and between the second and third trimesters. This study, which focused on immune-related processes in early gestation, was the first step in this investigation. The other limitation concerns the limited number of samples. The impact of both limitations is partly mitigated by comparing our results with available data obtained using microarray technology, which were not included in the study, and corresponding data obtained by RNA sequencing. We realize that data verification by classical RT-PCR, Western blot or histochemistry would be greatly appreciated.

## Conclusions

Our study is based on publicly available large-scale studies and their integrative analysis to enhance the statistical power of conclusions, mitigate the bias of various processing strategies, and consolidate and unify the accumulated results. The periodic integration and updating of large-scale data are necessary steps as new data accumulates.

**Table 1.** Differentially expressed genes in recognition and elimination of potential pathogen- or damage-associated molecular patterns.

| Gene symbol | Gene name | LogFC |
| --- | --- | --- |
| Pattern recognition receptors |  |  |
| *CD36* | Platelet glycoprotein 4 | 1.72 |
| *GPR34* | G protein-coupled receptor 34 | 1.55 |
| *GPR183* | G protein-coupled receptor 183 | 1.54 |
| *FPR1* | fMet-Leu-Phe receptor | 1.43 |
| *CD163* | Cluster of Differentiation 163 | 1.29 |
| *MRC1* | Mannose receptor C-type 1 | 1.17 |
| *TLR7* | Toll-like receptor 7 | 1.14 |
| *CD93* | Cluster of Differentiation 93 | 1.02 |
| Destruction of potential pathogens and damaged cells |  |  |
| *CYBB* | Cytochrome b-245 heavy chain (aka NOX2) | 1.47 |
| *RNASE6* | Ribonuclease K6 | 1.13 |
| *NCF2* | Neutrophil cytosolic factor 2 | 1.10 |
| *NCF4* | Neutrophil cytosolic factor 4 | 1.03 |
| *BST2* | Bone marrow stromal antigen 2 | 1.01 |
| *LCN2* | Neutrophil gelatinase-associated lipocalin | -1.98 |
| *SLPI* | Secretory leukocyte protease inhibitor | -2.18 |
| *DEFB1* | Beta-defensin 1 | -3.23 |

## Acknowledgements

This study was supported by the OeAD grant (# UA 06/2023).

## Author contributions (following the CReDiT Taxonomy)

MO headed conceptualization, writing of the original draft, project administration, supervision, and was involved in funding acquisition. OL contributed to conceptualization, data curation, formal analysis, methodology, database development, and validation. YK contributed to data deconvolution. OM contributed to data curation, formal analysis, methodology, and validation. VM contributed to GSEA analysis. YP contributed to updating the codebase and deploying the web interface of the database to the IGEA web portal. BH contributed to conceptualization and funding acquisition, and headed reviewing and editing of the draft and revised/figures.

## Data availability

Data for the analysis was obtained from the Gene Expression Omnibus (GEO) and BioStudies (ArrayExpress) databases. We developed the specialized IGEA (integrative gene expression analysis) database [https://molecularbiology.ink/igea/] containing the metadata for the genome-wide transcriptome profiling of more than 1000 samples of normal and pathology-affected human placentas. For direct inquiries, please contact the O. Lykhenko author. The entire pipeline code used in this study and the supplementary files are available via GitHub: https://github.com/Sashkow/r-affymetrix-integration-pipeline. It is a pipeline for Affymetrix microarray expression data integration, with an example of integrating 4 GEO datasets with samples of the first- and second-trimester human placentas.

## Document S1

**Table S1.** Datasets’ accession numbers (codes), sample information and references Table S1 is related to the main text, section 2, Methods

**Table S2.** Differentially expressed immune-related genes in the human placenta between the first and second trimesters.

Table S2 is related to the main text, section 3, Results and Discussion. Excel file containing additional data is too large to fit in a PDF. adj.P.Val – adjusted P-value.

**Table S3.** Differentially expressed genes in the migration of immune cells, cytokine activation, and immune tolerance.

Table S3 is related to the main text, section 3.2, Immune cell migration, cytokines, and immune tolerance

