## Supplementary material for "Placental immune factors change during the first half of healthy pregnancy": Document S1 (Tables S1, S3, S4)

**Table S1. Datasets’ accsession codes and sample information**

| Datasets  (GEО db) | І trimester | | ІІ trimester | | References |
| --- | --- | --- | --- | --- | --- |
|  | Samples,  number | Gestational  age, weeks | Samples,number | Gestational  age, weeks |  |
| GSE122214 | 4 | 7 - 8 |  |  | [1] |
| GSE22490 | 6 | 9 | 1 | 13 | [2] |
| GSE37901 |  |  | 4 | 17.1 - 18,8 | [3] |
| GSE9984 | 4 | 6.4 – 8.4 | 4 | 15.5 – 16.4 | [4] |
| Total number | 14 | | 9 | |  |

**Table S1 is related to the main text, section 2. Methods**

**Table S2. Differentially expressed immune-related genes between the second and first trimesters.**

**The table is related to the main text, section 3. Results and Discussion**

**Excel file containing additional data is too large to fit in a PDF.**

**adj.P.Val - adjusted P-value.**

**Table S3. Differentially expressed genes in the migration of immune cells,**

**cytokine activation, and immune tolerance**

| Gene symbol | Gene name | LogFC |
| --- | --- | --- |
| *Transendothelial cells migration* | | |
| *AOC1* | Diamine oxidase | 1.65 |
| *JAM2* | Junctional adhesion molecule B | 1.56 |
| *PVRL3* | Nectin-3 | 1.48 |
| *PECAM1* | Platelet endothelial cell adhesion molecule | 1.46 |
| *JAM-3* | Junctional adhesion molecule C | 1.09 |
| *EMCN* | Endomucin | **-**1.25 |
| *EpCAM* | Epithelial Cell Adhesion Molecule | **-**3.08 |
| *Chemokines, cytokines and receptors* | | |
| *CXCL7* | Pro-Platelet basic protein (PPBP) | 2.00 |
| *CXCL4* | Platelet factor 4 (PF4) | 1.98 |
| *EBI3* | Epstein-Barr virus-induced gene 3 | 1.89 |
| *S100A8* | Protein S100-A8 | 1.87 |
| *CXCL8* | Interleukin-8 | 1.74 |
| *S100-A9* | Protein S100-A9 | 1.70 |
| *IL-6* | Interleukin-6 | 1.69 |
| *CCL2* | C-C motif chemokine 2 | 1.65 |
| *CXCL1* | Growth-regulated alpha protein | 1.53 |
| *CCL13* | C-C motif chemokine 13 | 1.51 |
| *CSF2RB* | [Colony stimulating factor 2 receptor subunit beta](https://www.ncbi.nlm.nih.gov/gene/1439) | 1.44 |
| *CXCL2* | C-X-C motif chemokine 2 | 1.37 |
| *IL-12RВ2* | [Interleukin 12 receptor, beta 2 subuni](https://en.wikipedia.org/wiki/Interleukin_12_receptor,_beta_2_subunit)t | 1.25 |
| *CCL8* | C-C motif chemokine 8 | 1.14 |
| *SRGN* | Serglycin | 1.02 |
| *CXCL12* | C-X-C motif chemokine 12 | **-**1.17 |
| *CCL21* | C-C motif chemokine 21 | **-**1.24 |
| *CCR7* | C-C chemokine receptor type 7; | **-**1.26 |
| *CXCL13* | C-X-C motif chemokine 13 | **-**2.18 |
| *Immune tolerance and IgG transplacental transport* | | |
| *FCGR2B* | Fc fragment of IgG receptor IIb | 2.97 |
| *LAIR1* | Leukocyte associated immunoglobulin like receptor 1 | 1.34 |
| *LILRB* | Leukocyte immunoglobulin like receptor B5 | 1.27 |
| *PAEP* | Progestagen-associated endometrial protein | -4.43 |

**Table S3 Is related to the main text, section 3.2, Immune cell migration, cytokines, and immune tolerance**

**Table S4. The enriched biological processes in the differentially expressed genes between first and second trimesters of gestation**

| id | source | term_id | term_name | term_size | p.adj-value |
| --- | --- | --- | --- | --- | --- |
| 1 | GO:BP | GO:0000819 | sister chromatid segregation | 222 | 4.051786e-08 |
| 2 | GO:BP | GO:0000070 | mitotic sister chromatid segregation | 183 | 3.396686e-08 |
| 3 | GO:BP | GO:0140014 | mitotic nuclear division | 268 | 1.120974e-05 |
| 4 | GO:BP | GO:0007059 | chromosome segregation | 404 | 9.629898e-07 |
| 5 | GO:BP | GO:0098813 | nuclear chromosome segregation | 304 | 3.027122e-07 |
| 6 | GO:BP | GO:0051276 | chromosome organization | 600 | 1.817684e-06 |
| 7 | GO:BP | GO:0048285 | organelle fission | 465 | 6.506322e-06 |
| 8 | GO:BP | GO:0000280 | nuclear division | 420 | 1.214271e-06 |
| 9 | GO:BP | GO:1903047 | mitotic cell cycle process | 711 | 7.002168e-06 |
| 10 | GO:BP | GO:0051301 | cell division | 613 | 1.394302e-07 |
| 11 | GO:BP | GO:0051303 | establishment of chromosome localization | 109 | 9.128471e-06 |
| 12 | GO:BP | GO:0050000 | chromosome localization | 117 | 5.790280e-06 |
| 13 | GO:BP | GO:0007094 | mitotic spindle assembly checkpoint signaling | 46 | 1.044370e-05 |
| 14 | GO:BP | GO:0031577 | spindle checkpoint signaling | 47 | 9.447887e-06 |
| 15 | GO:BP | GO:0071173 | spindle assembly checkpoint signaling | 46 | 1.044370e-05 |
| 16 | GO:BP | GO:0071174 | mitotic spindle checkpoint signaling | 46 | 1.044370e-05 |
| 17 | GO:BP | GO:0007091 | metaphase/anaphase transition of mitotic cell cycle | 92 | 1.067064e-05 |
| 18 | GO:BP | GO:0040015 | negative regulation of multicellular organism growth | 12 | 9.749256e-06 |
| 19 | GO:BP | GO:0048771 | tissue remodeling | 168 | 1.441014e-06 |
| 20 | GO:BP | GO:0001525 | angiogenesis | 519 | 8.326415e-06 |
| 21 | GO:BP | GO:0048545 | response to steroid hormone | 326 | 4.471611e-07 |
| 22 | GO:BP | GO:0050886 | endocrine process | 91 | 5.651933e-06 |
| 23 | GO:BP | GO:0060986 | endocrine hormone secretion | 55 | 4.035228e-06 |
| 24 | GO:BP | GO:0070848 | response to growth factor | 684 | 2.177534e-06 |
| 25 | GO:BP | GO:0071363 | cellular response to growth factor stimulus | 653 | 1.041790e-06 |
| 26 | GO:BP | GO:0009991 | response to extracellular stimulus | 499 | 1.774747e-06 |
| 27 | GO:BP | GO:0031667 | response to nutrient levels | 470 | 6.677526e-07 |
| 28 | GO:BP | GO:0044703 | multi-organism reproductive process | 189 | 1.801798e-06 |
| 29 | GO:BP | GO:0044706 | multi-multicellular organism process | 196 | 2.715381e-07 |
| 30 | GO:BP | GO:0007565 | female pregnancy | 170 | 8.452893e-07 |

**Table S4 is related to the main text, section 3.3, Placental cell type deconvolution.**
